# The thalamic reticular nucleus-lateral habenula circuit regulates depressive-like behaviors in chronic stress and chronic pain

**DOI:** 10.1101/2023.02.17.528253

**Authors:** Xiang Xu, Rui Chen, Xin-Yue Wang, Wen-Bin Jia, Peng-Fei Xu, Xiao-Qing Liu, Ying Zhang, Xin-Feng Liu, Yan Zhang

## Abstract

Chronic stress and chronic pain are two major predisposing factors to trigger depression. Enhanced excitatory input to the lateral habenula (LHb) has been implicated in the pathophysiology of depression. However, the contribution of inhibitory transmission remains elusive. Here, we dissect an inhibitory projection from the sensory thalamic reticular nucleus (sTRN) to LHb, which is activated by acute aversive stimuli. However, chronic restraint stress (CRS) weakens sTRN-LHb synaptic strength, and this synaptic attenuation is indispensable for CRS-induced LHb neural hyperactivity and depression onset. Moreover, artificially inhibiting sTRN-LHb circuit induces depressive-like behaviors in healthy mice, while enhancing this circuit relieves depression induced by both chronic stress and chronic pain. Intriguingly, neither neuropathic pain nor comorbid pain in chronic stress is affected by this pathway. Together, our study demonstrates a novel sTRN-LHb circuit in establishing and modulating depression, thus shedding light on potential therapeutic targets for preventing or managing depression.

## Introduction

Depression is a common mood disorder and a leading cause of disability around the world. Chronic stress and chronic pain are the two most common contributors that can lead to psychological dysfunctions such as depression. Alternatively, depressed patients have a high incidence of pain (*1*). The vicious cycle of depression-pain comorbidity brings a great challenge for refractory depression management (*2*-*4*).

The LHb is an evolutionarily conserved epithalamic nucleus in vertebrates (*5*). As a primary negative regulator of monoaminergic brain regions, the LHb has been implicated in encoding negative outcomes and aversive behaviors (*6*, *7*). Preclinical studies revealed that the glutamatergic neurons of LHb (referred to as LHb^GLU^ neurons) can be activated instantly by aversive events (*8*, *9*). After exposure to chronic restraint stress or chronic pain, the mice displayed hyperactivity of the LHb^GLU^ neurons and depressive-like behaviors (*9*-*11*). On the contrary, LHb lesions or LHb neuronal suppression improves depressive-like symptoms in rodents (*9*, *10*, *12*).

Clinically, LHb neuronal activity is increased in patients with depression (*13*), and deep brain stimulation to inactivate LHb has been used to relieve major depression (*14*). These discoveries suggest the compelling association of LHb dysfunction with depression. The potential role of LHb in the processing of pain and analgesic signals has also been reported (*15*-*17*).

The maladaptive neuronal dysfunction could arise from changes in intrinsic properties or synaptic changes caused by the imbalance of presynaptic GABAergic and glutamatergic transmission (*18*). The LHb receives extensive excitatory inputs from the limbic forebrain regions and basal ganglia (*7*). Hyperactivity of the LHb^GLU^ neurons has been linked to enhanced excitatory inputs (*9*, *19*, *20*). Manipulation of LHb-upstream excitatory afferents, such as lateral hypothalamus (LH) (*9*, *21*), substantia innominate (*8*), medial prefrontal cortex (mPFC) (*22*), lateral preoptic areas (LPO) (*23*) and ventral pallidum (VP) (*24*), can bidirectionally regulate depressive-like behaviors. However, much less attention has been paid to the role of inhibitory afferents to LHb in the pathophysiology and modulation of depression.

The thalamic reticular nucleus (TRN), a cluster of GABAergic neurons (*25*), is a thin, shell-like structure located between the dorsal thalamus and cerebral cortex. It receives inputs from the cortex and other thalamic nuclei. Meanwhile, the TRN provides the major inhibition to thalamic neurons and functions as a “gateway filter” in information flow between the cortex and dorsal thalamus (*26*-*28*). The TRN has been elucidated in arousal, cognitive function, sensorimotor processing, defensiveness, and pain modulation (*29*-*34*), yet the functional role of TRN-LHb projection in depression and pain has not been explored.

Here, we found that the somatostatin-expressing neurons, rather than parvalbumin (PV)-expressing neurons in sensory TRN (referred to as sTRN^SOM^ and sTRN^PV^ neurons, respectively), send direct inhibitory inputs to LHb^GLU^ neurons which are involved in both aversive information and pain processing. *In vivo* fiber photometry revealed that the TRN afferents in the LHb are instantly activated by acute aversive stimuli. *In vitro* electrophysiological recordings further demonstrated that sTRN^SOM^-LHb synaptic connectivity is blunted by CRS. Notably, repetitive activation of the sTRN-LHb circuit is sufficient to prevent depression formation by altering the excitability of LHb neurons in mice subjected to CRS. Furthermore, artificial manipulation of this circuit bidirectionally modulates depressive-like behaviors, rather than pain-like behaviors. Finally, whole-brain mapping dissects the brain regions which are involved in signaling stress information to sTRN. Thus, this study systematically elucidates a novel inhibitory sTRN-LHb circuit in the pathophysiology of depression which provides a functional substrate for early intervention before depression onset and also for depression management.

## Results

### Aversive stimuli activate LHb-projecting sTRN neurons

Somatostatin-expressing and pavalbumin-expresssing neurons are two major subsets of inhibitory neurons within the TRN (*35*). To assess the TRN-LHb projection, we injected anterograde tracing virus AAV-DIO-mCherry into the caudal part (also referred to as sTRN) or rostral part (also referred to as lTRN) of TRN in *SOM-Cre* and *PV-Cre* mice (Fig. 1A and fig. S1, A and C). Intriguingly, dense mCherry^+^ fibers were detected exclusively in LHb of *SOM-Cre* mice with sTRN injection (Fig. 1, A-C and fig. S1, A-D). To exclude the passenger fibers across the LHb, the sTRN^SOM^-LHb projection was further supported by retrograde trans-monosynaptic labeling by injecting AAV2/2-retro-hSyn-Cre into the LHb and AAV-DIO-mCherry into the sTRN of *C57* mice (Fig. 1, D and E), showing that ∼89% of retrogradely labeled mCherry^+^ neurons in sTRN overlapped with somatostatin. Furthermore, AAV-based anterograde trans-monosynaptic labeling in *C57* mice proved that sTRN-targeted LHb neurons were major (∼97%) glutamatergic neurons (Fig. 1, F and G). From the data, we proved that sTRN^SOM^ neurons send direct projections to LHb^GLU^ neurons.

**Fig. 1.**
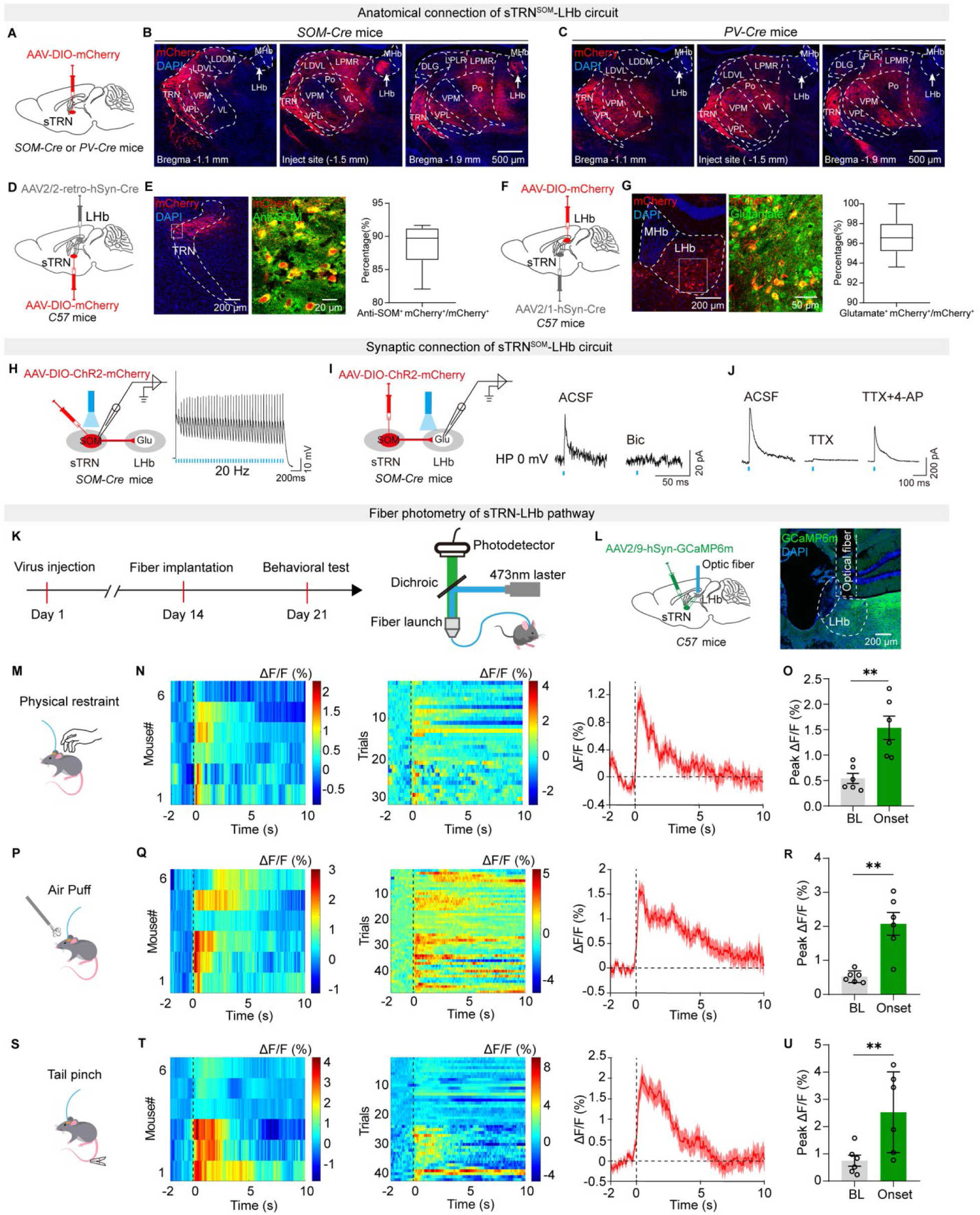
Aversive stimuli activate LHb-projecting sTRN afferents. (A) Schematic showing viral injection into the sensory TRN of *SOM-Cre* or *PV-Cre* mice. (B and C) Representative images of mCherry-expressing signals in different brain regions of *SOM-Cre* (B) or *PV-Cre* (C) mice. Scale bars, 500 μm. (D and E) Schematic of the Cre-dependent retrograde trans-monosynaptic tracing strategy in *C57* mice (D) and summary data for the percentage of mCherry^+^ neurons in the sTRN which co-localize with anti-SOM immunofluorescence (E). Scale bars, 200 μm and 20 μm. (n=6 sections from three mice) (F and G) Schematic of the Cre-dependent anterograde trans-monosynaptic tracing strategy in *C57* mice (F) and summary data for the percentage of mCherry^+^ neurons in the LHb which co-localize with anti-Glutamate immunofluorescence (G). Scale bars, 200 μm and 50 μm. (n=10 sections from five mice) (H) Schematic showing sTRN electrophysiological recordings in acute slices from *SOM-Cre* mice (left) and a representative trace of blue light (473 nm, 20 Hz)-evoked action potentials in a ChR2-mCherry-expressing sTRN neuron (right). (I) Schematic showing LHb electrophysiological recordings in acute slices from *SOM-Cre* mice (left) and representative traces of light-evoked IPSCs of sTRN neurons before (ACSF) and after bicuculline (Bic, 10 μM) treatment. (J) Representative traces of light-evoked IPSCs of the LHb neurons before (ACSF) and after TTX (1 μM) or TTX and 4-AP (500 μM) treatment. (K) Schematic of experimental design (left) and fiber photometry recording *in vivo* (right). (L) Illustration (left) and representative image (right) of viral delivery and optic fiber implantation. Scale bar, 200 μm. (M, P and S) Schematic of fiber photometry recording in response to RS (M), air puff (P) and tail pinch (S). (N, Q and T) Heatmap and average responses showing Ca^2+^ transients evoked by RS (N), air puff (Q), and tail pinch (T) in the LHb neurons. (n=6 mice/group) (O, R and U) Quantification of peak average Ca^2+^ responses before and after RS (O), air puff (R), and tail pinch (U) stimulation (right). (n=6 mice/group) For (E) and (G), data are shown as box and whisker plots (medians, quartiles (boxes) and ranges minimum to maximum (whiskers)); For (O), (R), and (U), data are presented as mean ± SEM. “ns”, no significance; **p < 0.01. Paired two-sided t test for (O), (R), and (U).

To examine functional projections from the sTRN to the LHb, AAV-DIO-ChR2-mCherry was injected into the sTRN of *SOM-Cre* mice (Fig. 1H, left). The functional viral expression was proved by the reliable action potentials of ChR2-expressing sTRN^SOM^ neurons following 20 Hz blue light stimulation (Fig. 1H, right). Whole-cell patch-clamp recordings from the LHb neurons showed that photostimulation of ChR2^+^ fibers originating in the sTRN evoked inhibitory postsynaptic currents (IPSCs), which can be blocked by bath application of bicuculline (Bic; 10 μM), a selective GABA_A_ receptor antagonist (Fig. 1I). Moreover, light-evoked IPSC was blocked by tetrodotoxin (TTX) and partially restored by potassium channel blocker 4-aminopyridine (4-AP) (Fig. 1J), suggesting that LHb receives direct inhibitory inputs from the sTRN^SOM^ neurons.

The LHb is perceived as a key hub for aversive information processing. Consistent with previous studies (*8*, *9*), LHb neurons were activated in mice subjected to RS (6 h), compared with mice with fasting for solids and liquids (6 h) but remained unrestraint (fig. S2, A and B). *In vivo* fiber photometry further demonstrated that population activity of LHb^GLU^ neurons was significantly increased upon various aversive stimuli including acute RS and tail pinch (fig. S2, C-H and L-N). A strong tendency of calcium signal increment was observed in air puff assay (fig. S2, I-K). We next want to know whether the sTRN-LHb projection also processes aversive information. We recorded Ca^2+^ transients of sTRN afferents within LHb (Fig. 1K). To this end, we expressed Ca^2+^ indicator GCaMP6m in sTRN and implanted an optical fiber above the LHb of *C57* mice (Fig. 1L). We observed bulk calcium signals of LHb-projecting sTRN afferents in response to acute RS, air puff, or tail pinch (Fig. 1, M-U), suggesting that sTRN-LHb projection is also behaviorally engaged in processing aversive information.

### Chronic restraint stress (CRS) drives sTRN-LHb synaptic attenuation

The balance between GABAergic and glutamatergic transmission controls LHb activity and behavior (*18*). Enhanced excitatory transmission to LHb has been reported in different animal models of depression (*9*, *19*, *20*). However, whether the inhibitory transmission of the sTRN-LHb circuit would be affected by chronic stress, one of the most common factors causing depression onset, is unknown.

To resolve the question, we assessed the synaptic efficacy of the sTRN-LHb circuit after chronic restraint stress (CRS, 3 weeks restraint for 6 hours per day). As reported previously (*10*, *36*), CRS induced depressive-like behaviors, including the increased immobile duration in both the forced swimming test (FST) and tail suspension test (TST), and decreased sucrose preference ratio in sucrose preference test (SPT) (fig. S3, A and B). The locomotion tested in the open-field test (OFT) was not affected (fig. S3C).

To evaluate the synaptic transmission of the sTRN-LHb circuit, we injected Cre-dependent AAV-DIO-ChR2-mCherry into the sTRN of *SOM-Cre* mice and performed whole-cell recordings of LHb neurons in slices obtained from CRS and control mice (Fig. 2A). We found that the amplitude of light-evoked IPSCs was significantly decreased in CRS mice compared with naïve controls (Fig. 2, B and C). We next assessed paired-pulse ratio (PPR) of light-evoked IPSCs, which is inversely correlated with the presynaptic transmitter release (Fig. 2, B and D). We found that paired-pulse ratio (PPR) of the sTRN^SOM^-LHb synapse was significantly increased in CRS mice compared with naïve controls (Fig. 2D), implying presynaptic mechanism involvement. Consistently, the frequency of miniature IPSCs (mIPSCs) was reduced in CRS mice compared with naïve controls (Fig. 2, E and F). Meanwhile, the amplitude of mIPSCs was also attenuated (Fig. 2, E and G). These data suggest that CRS blunts the synaptic strength of the sTRN^SOM^-LHb circuit via attenuation of both the presynaptic GABA release and postsynaptic GABA receptor activity.

**Fig. 2.**
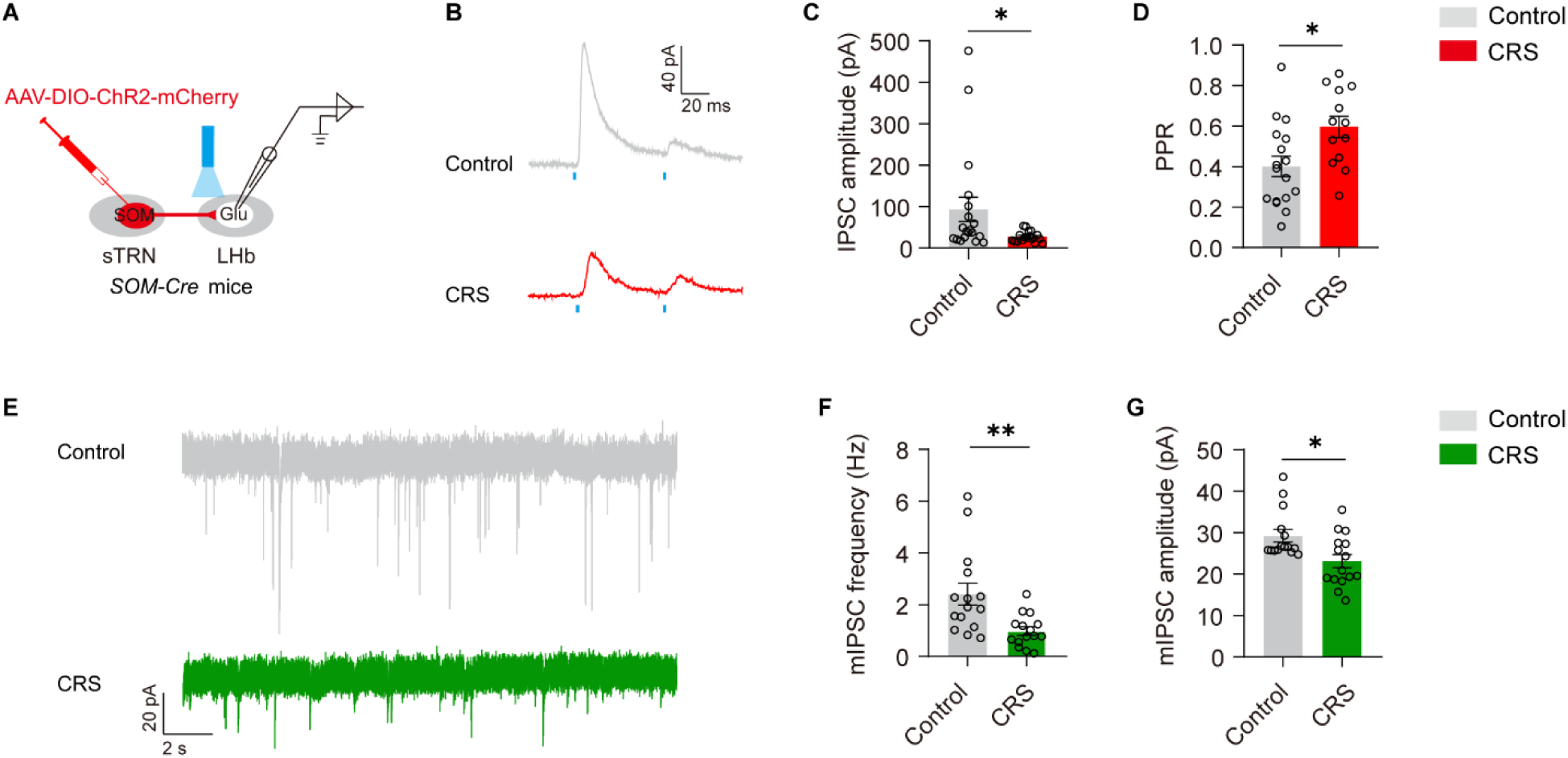
Chronic restraint stress (CRS) drives sTRN^SOM^-LHb synaptic attenuation. (A) Schematic showing LHb electrophysiological recordings in acute slices. (B) Representative traces of light-evoked paired-pulse ratio (PPR) recorded from LHb neurons of control or CRS mice. (C and D) Summary data for the amplitude (C) and PPR (D) of light-evoked IPSCs recorded from the sTRN^SOM^-targeted LHb neurons of control or CRS mice. (E) Representative traces of mIPSC recorded from LHb neurons of control or CRS mice. (F and G) Summary data for the frequency (F) and amplitude (G) of mIPSC recorded from the LHb neurons of control or CRS mice. Data are presented as mean ± SEM. *p < 0.05, **p < 0.01. Mann-Whitney U test for (C), (F), and (G); Unpaired two-sided t test for (D).

### Repeated activation of the LHb-projecting sTRN neurons prevents CRS-induced depression onset by altering the excitability of LHb neurons

Considering that sTRN-LHb synaptic efficacy was undermined under CRS, we went on to explore whether artificially enhancing LHb-projecting sTRN neural activity during CRS could prevent depression onset. We thus injected AAV2/2-retro-hSyn-Cre into the LHb and AAV-DIO-hM3D(Gq)-mCherry or AAV-DIO-mCherry into the sTRN of *C57* mice to activate the LHb upstream sTRN neurons during CRS (Fig. 3, A and B). The efficacy of virus expression was confirmed by depolarization of the resting membrane potentials (RMP) and neural firing by CNO in hM3Dq-expressing sTRN neurons (Fig. 3C). We then intraperitoneally injected CNO before daily RS (6 h) for 21 days. Interestingly, we found that chronic excitation of the LHb-projecting sTRN neurons during CRS successfully prevented the development of depression in hM3Dq group, while the mCherry control mice exhibited depressive-like behaviors assessed by the FST, TST, and SPT (Fig. 3, D-F). The locomotion assessed by OFT was not affected by repeated activation of the LHb-projecting sTRN neurons (Fig. 3G).

**Fig. 3.**
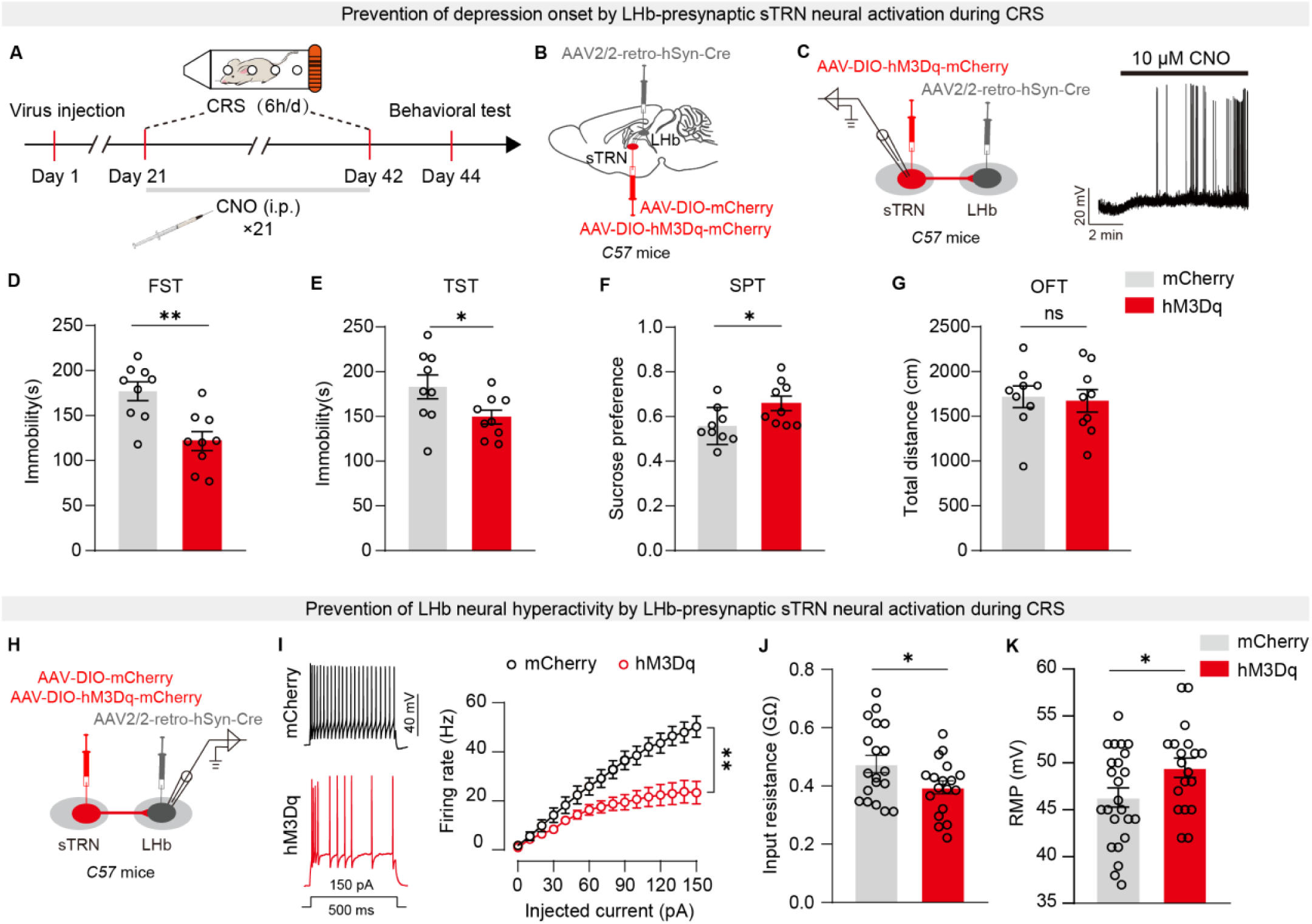
Repeated activation of the LHb-projecting sTRN neurons prevents CRS-induced depression onset by altering the excitability of LHb neurons. (A) Schematic of experimental design. (B) Illustration of viral delivery. (C) Schematic showing sTRN electrophysiological recordings in acute slices (left) and a representative trace showing depolarization of the hM3Dq-mCherry-expressing sTRN neuron by CNO (10 μM). (D-G) Behavioral effects of repeated activation of the LHb-projecting sTRN neurons during CRS on depressive-like behaviors assessed by FST (D), TST (E), and SPT (F), and locomotion activity assessed by OFT (G) in *C57* mice. (n=9 mice/group) (H) Schematic showing LHb electrophysiological recordings in acute slices. (I-K) Summary data for the firing rate (I), input resistance (J), and resting membrane potential (K) recorded from LHb neurons of mCherry&CRS or hM3Dq&CRS mice. Data are presented as mean ± SEM. “ns”, no significance; *p < 0.05, **p < 0.01. Unpaired two-sided t test for (D-G), (J), and (K); Two-way repeated measures ANOVA with Bonferroni *post hoc* analysis for (I).

Given that hyperactivity of LHb neurons is closely associated with the pathophysiology of depression (*9*, *19*, *20*, *37*), we further explored whether chemogenetic excitation of the LHb-projecting sTRN neurons would prevent the hyperactivity of LHb neurons induced by CRS. First of all, we found that the firing rate of LHb neurons was significantly increased in slices obtained from CRS mice compared with naïve controls (fig. S3, D and E). Meanwhile, the LHb neurons displayed depolarized resting membrane potential (RMP), and increased input resistance in slices obtained from CRS mice compared with naïve controls (fig. S3, F and G). However, chronic activation of the LHb-projecting sTRN neurons during CRS restored the hyperexcitablity of LHb neurons, as indicated by the decreased neural firing frequency, hyperpolarized RMP and decreased input resistance of LHb neurons in CRS-treated hM3Dq mice compared with CRS-treated mCherry controls (Fig. 3, H-K). Taken together, attenuated sTRN inputs are indispensable for CRS-induced hyperactivity of LHb neurons and depression onset.

### Acute inhibition of the sTRN^SOM^-LHb circuit induces depressive-like behaviors in naïve mice

The causal link between attenuated sTRN^SOM^-LHb synaptic strength with CRS-induced depression prompted us to further examine whether acute inhibition of the sTRN^SOM^-LHb circuit would induce depressive-like behaviors in naïve mice. To this end, we injected AAV-DIO-eNpHR3.0-EYFP or AAV-DIO-EYFP into the sTRN and implanted optical fiber above the LHb of *SOM-Cre* mice (Fig. 4, A-C). Current clamp recordings on eNpHR3.0-expressing sTRN^SOM^ neurons showed that 20 Hz yellow light stimulation suppressed action potentials (Fig. 4D), suggesting functional viral transduction. Behaviorally, optogenetic inhibition of the sTRN^SOM^ afferents within LHb was sufficient to induce depressive-like behaviors assessed by the TST and SPT (Fig. 4, E and F), which is similar to depressive-like behaviors induction following chemogenetic activation of LHb^GLU^ neurons (fig. S3, J-L). The locomotion tested in OFT was unaffected by the sTRN^SOM^-LHb inhibition or LHb^GLU^ neural activation (Fig. 4G and fig. S3M). These data indicate that tonic activity of sTRN^SOM^ afferents within LHb is necessary for preventing depressive-like behaviors in naïve mice.

**Fig. 4.**
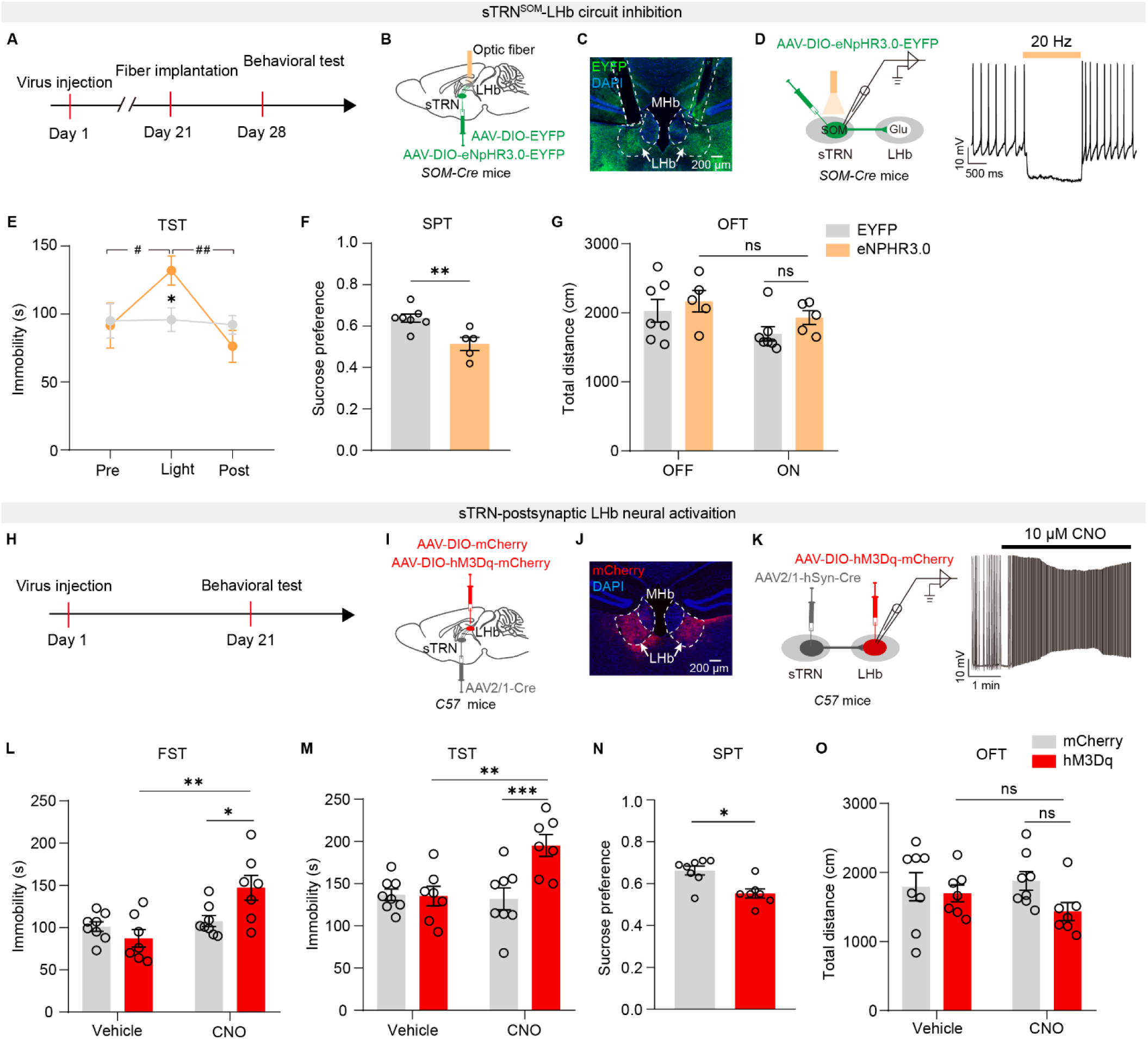
Acute inhibition of the sTRN^SOM^-LHb circuit induces depressive-like behaviors in naïve mice. (A and H) Schematic of experimental design. (B and C) Illustration (B) and representative image (C) of viral delivery and bilateral optic fibers implantation. Scale bar, 200 μm. (D) Schematic of recording configuration in acute slices and representative trace showing suppression of action potentials in an eNpHR3.0-EYFP^+^ sTRN neuron by laser stimulation (584 nm, 20 Hz). (E-G) Behavioral effects of optogenetic inhibition of sTRN^SOM^-LHb circuit on depressive-like behaviors assessed by TST (E) and SPT (F), and locomotion activity assessed by OFT (G) in naïve *SOM-Cre* mice. (n=5-7 mice/group) (I and J) Illustration (I) and representative image (J) of viral delivery. Scale bar, 200 μm. (K) Schematic of recording configuration in acute slices and representative trace showing depolarization of membrane potential in a hM3Dq-mCherry^+^ LHb neuron by CNO (10 μM). (L-O) Behavioral effects of chemogenetic activation of sTRN-postsynaptic LHb neurons on depressive-like behaviors assessed by FST (L), TST (M), and SPT (N), and locomotion activity assessed by OFT (O) in naïve *C57* mice. (n=7-8 mice/group) Data are presented as mean ± SEM. “ns”, no significance; *p < 0.05, **p < 0.01, ***p < 0.001; ^#^p < 0.05, ^##^p < 0.01. Two-way repeated measures ANOVA with Bonferroni *post hoc* analysis for (E), (G), (L), (M), and (O); Unpaired two-sided t test for (F); Mann-Whitney U test for (N).

To further demonstrate the role of the sTRN-LHb circuit in depression, we also investigated the sufficiency of sTRN-postsynaptic LHb neural activation in driving depressive-like behaviors. We injected the AAV2/1-hSyn-Cre into the bilateral sTRN and AAV-DIO-hM3D(Gq)-mCherry or AAV-DIO-mCherry into the LHb of *C57* mice (Fig. 4, H-J). After 21 days of viral expression, the sTRN-targeted LHb neurons expressed functional hM3Dq indicated by neural depolarization by CNO (10 μM) application (Fig. 4K). The mice were then subjected to a series of depressive-like behavioral tests following i.p CNO or vehicle. We found that activation of sTRN-postsynaptic LHb neurons was also sufficient to drive depressive-like behaviors tested in FST, TST, and SPT (Fig. 4, L-N), without any influence on locomotion (Fig. 4O). In summary, sTRN^SOM^-LHb circuit inhibition induces depressive-like behaviors.

### Activation of the sTRN^SOM^-LHb circuit alleviates depressive-like behaviors induced by chronic stress and chronic pain

Given that LHb neural hyperactivity underlies depression and sTRN^SOM^-LHb projections are GABAergic, we postulated that activation of the sTRN^SOM^-LHb circuit would alleviate depression. Because chronic stress and chronic pain are the two most common contributors causing depression, we thus tested the postulation on both CRS-induced depression and neuropathic pain-induced comorbid depression models.

First of all, the CRS-induced depression model was studied. We injected *SOM-Cre* mice with AAV-DIO-ChR2-mCherry or AAV-DIO-mCherry into the LHb and subsequently performed CRS for 21 days (Fig. 5, A-C). After that, optical fibers were implanted above the LHb, and mice were allowed for an additional one-week recovery before subjecting to depression-related behavioral tests. As expected, optogenetic activation of sTRN^SOM^ neural terminals within the LHb significantly rescued depressive-like behaviors tested in the TST and SPT in the ChR2 group compared with mCherry controls (Fig. 5, D and E), which is similar to depressive-like behaviors relief induced by chemogenetic inhibition of LHb^GLU^ neurons (fig. S4, A-E). The locomotion tested in OFT was unaffected by the circuit inhibition or LHb^GLU^ neural activation (Fig. 5F and fig. S4F).

**Fig. 5.**
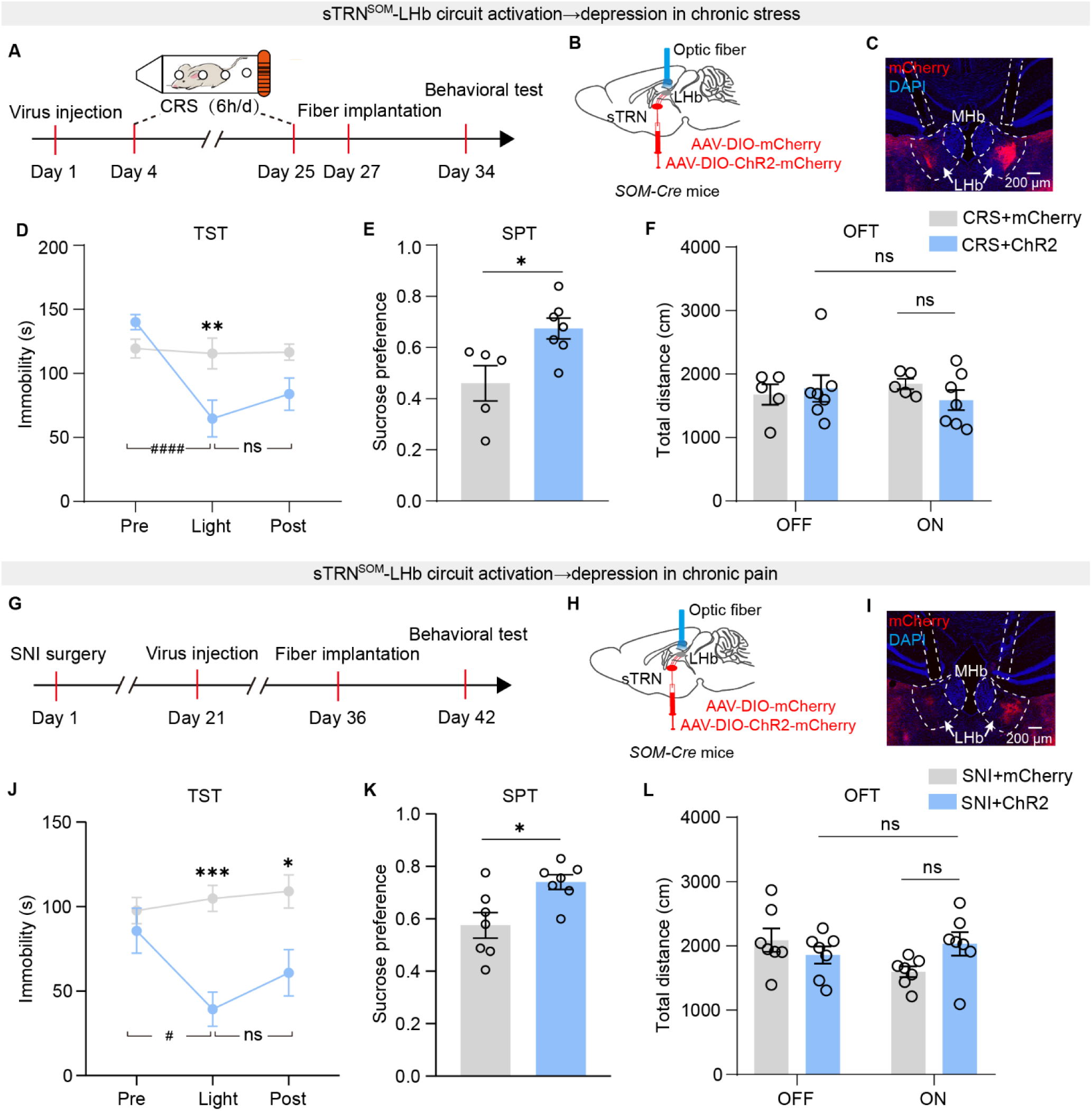
Activation of the sTRN^SOM^-LHb circuit alleviates depressive-like behaviors induced by chronic stress and chronic pain. (A and G) Schematic of experimental design. (B and C) Illustration (B) and representative image (C) of viral delivery and bilateral optic fibers implantation. Scale bar, 200 μm. (D-F) Behavioral effects of optogenetic activation of LHb-projecting sTRN^SOM^ neural afferents on depressive-like behaviors assessed by TST (D) and SPT (E), and locomotion activity assessed by OFT (F) in CRS-treated *SOM-Cre* mice. (n=5-7 mice/group) (H and I) Illustration (H) and representative image (I) of viral delivery and bilateral optic fibers implantation. Scale bar, 200 μm. (J-L) Behavioral effects of optogenetic activation of LHb-projecting sTRN^SOM^ neural afferents on depressive-like behaviors assessed by TST (J) and SPT (K), and locomotion activity assessed by OFT (L) in SNI-treated *SOM-Cre* mice. (n=7 mice/group) Data are presented as mean ± SEM. “ns”, no significance; *p < 0.05, ***p < 0.001; ^#^p < 0.05, ^####^p < 0.0001. Two-way repeated measures ANOVA with Bonferroni *post hoc* analysis for (D), (F), (J) and (L); Unpaired two-sided t test for (E) and (K).

To further investigate the role of the sTRN-LHb circuit in depression relief, we examined whether sTRN-postsynaptic LHb neural inhibition could relieve CRS-induced depressive-like behaviors. We bilaterally injected the AAV2/1-hSyn-Cre into sTRN and AAV-DIO-hM4D(Gi)-mCherry or AAV-DIO-mCherry into LHb of *C57* mice, which were then restrained for 21 days (fig. S4, G-I). Patch-clamp recording from hM4Di-expressing neurons in the LHb showed the hyperpolarization of the resting membrane potential by CNO (fig. S4J), confirming the functionality of the virus. After i.p CNO, CRS-induced depressive-like behaviors were largely alleviated in the hM4Di group compared with saline-treated and mCherry-injected naïve controls (fig. S4, K-M). The motor activity assessed by OFT was not affected by sTRN-targeted LHb neural inhibition (fig. S4N).

Second, we investigated the role of the sTRN-LHb circuit in comorbid depression induced by chronic pain, which is one of the most common forms of drug-resistant depression. Here, we used spared nerve injury (SNI) as a chronic pain model (Fig. 5G). Six weeks after the operation, mice developed depressive-like symptoms in the FST, TST, and SPT assays (fig. S5, A and B), without any deficit in locomotion activity (fig. S5C). Chemogenetic inhibition of LHb^GLU^ neurons can greatly relieve neuropathic pain-induced comorbid depressive-like behaviors (fig. S5, D-G) without any influence on locomotion (fig. S5H), suggesting the involvement of LHb in the comorbid depression under chronic pain as previously reported (*11*). Moreover, either optogenetic activation of sTRN^SOM^ neural afferents within the LHb (Fig. 5, J and K) or the specific inhibition of sTRN-targeted LHb neurons (fig. S5, I-L) can efficiently alleviate comorbid depressive-like behaviors in neuropathic pain, as observed in CRS model. The locomotion was unaltered by both manipulations (Fig. 5L and Fig. S5M). Taken together, enhancing the sTRN^SOM^-LHb circuit can rescue depressive-like behaviors induced by both chronic stress and chronic pain.

### Activation of the sTRN^SOM^-LHb circuit does not affect pain induced by nerve injury and depression

Besides the pivotal role in depression, the LHb was also involved in pain processing (*15*-*17*). We next investigated whether sTRN^SOM^-LHb circuit activation would affect pain in two models: SNI-induced neuropathic pain and comorbid pain in depression. Surprisingly, sTRN^SOM^-LHb circuit activation did not relieve the SNI-induced mechanical hypersensitivity (Fig. 6, A-C). Given the high incidence of comorbid pain in depressive patients, we repeated the experiment as performed in the SNI model. As previous study (*36*), the mice displayed marked mechanical hypersensitivity until three weeks CRS of when depressive-like behaviors developed, and the comorbid pain lasted at least two weeks after termination of CRS (Fig. 6, D and E). Next, we evaluated the effect of sTRN^SOM^-LHb circuit activation on pain threshold within the two weeks-time windows. We observed that mechanical hypersensitivity induced by CRS was also unaffected (Fig. 6F). Collectively, our data indicate that the sTRN^SOM^-LHb circuit specifically modulates depression rather than pain.

**Fig. 6.**
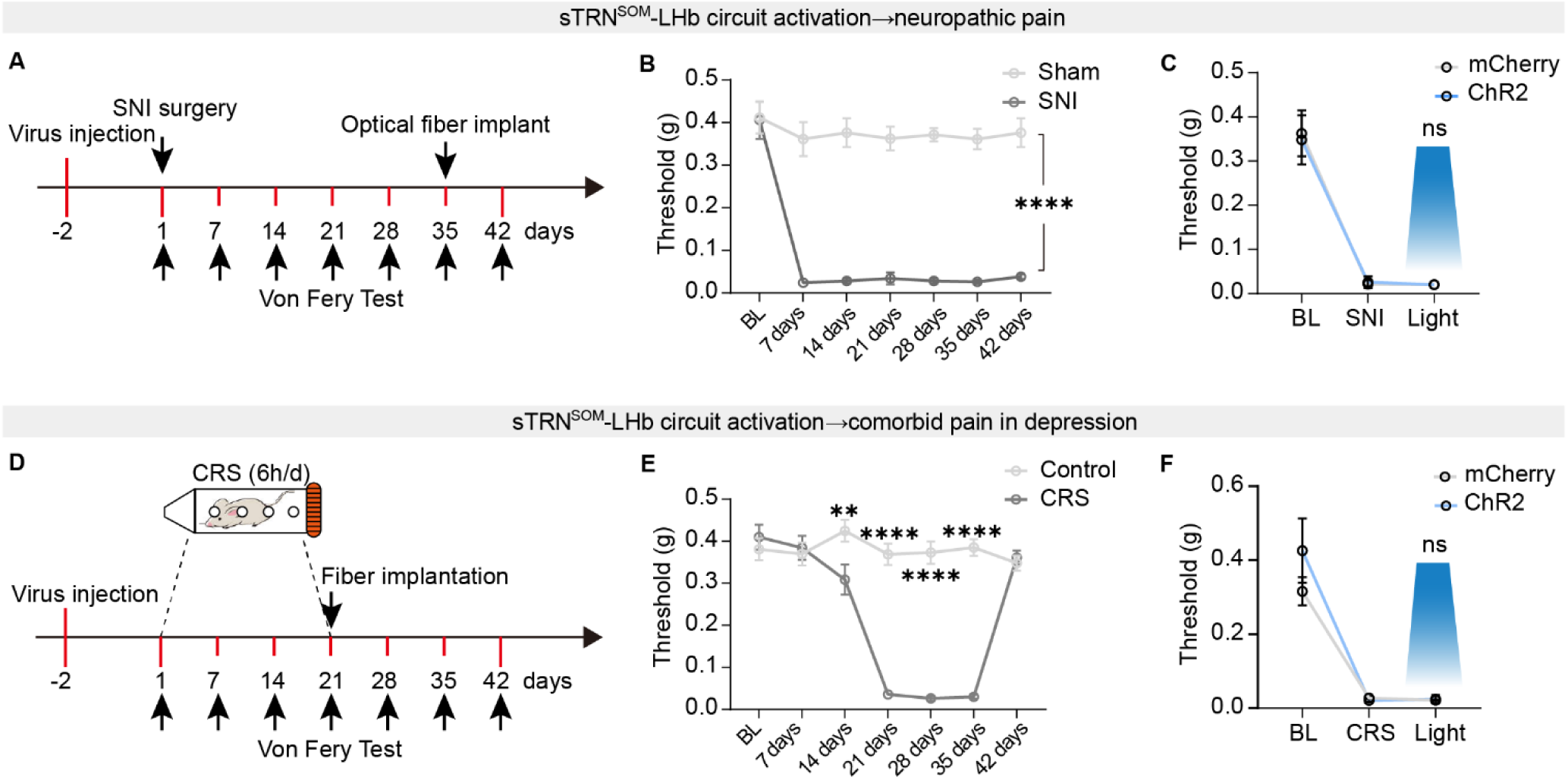
Activation of the sTRN^SOM^-LHb circuit does not affect pain induced by nerve injury and depression. (A and D) Schematic of experimental design. (B) Mechanical hypersensitivity in SNI-treated *C57* mice. (n=5-8 mice/group) (C) Behavioral effects of optogenetic activation of LHb-projecting sTRN^SOM^ neural afferents on pain-like behavior assessed by von Frey test in SNI-treated *SOM-Cre* mice. (n=5-7 mice/group) (E) Comorbid pain-like behavior in CRS-treated *C57* mice. (n=7-10 mice/group) (F) Behavioral effects of optogenetic activation of LHb-projecting sTRN^SOM^ neural afferents on pain-like behaviors assessed by von Frey test in CRS-treated *SOM-Cre* mice. (n=5-7 mice/group) Data are presented as mean ± SEM. “ns”, no significance; **p < 0.01, ****p < 0.0001. Two-way repeated measures ANOVA with Bonferroni *post hoc* analysis for (B), (C), (E) and (F).

### sTRN upstream brain regions associated with stress

As early as 1984, the TRN was described as the “guardian of the gateway” in the thalamocortical circuit (*27*). We next investigate its upstream brain regions that are associated with stress encoding. To do this, AAV2/2-retro-hSyn-Cre was injected into the sTRN of *Ai14* mice (fig. S6A). Three weeks later when sTRN upstream brain areas were labeled, mice were further exposed to RS (6h) or remained undisturbed (fig. S6A). We found that sTRN received broad presynaptic inputs from various brain regions (fig. S6, B-I), including the central amygdala (CeA), basolateral amygdala (BLA), ventral medial nucleus (VM), anterior cingulate cortex (ACC), insular cortex (IC) and dorsal raphe nucleus (DRN). Within these brain areas, we observed many c-Fos-positive neurons in mice exposed to 6h RS while rare signals were detected in undisturbed controls (fig. S6, C and E-I). Moreover, a small percentage of tdTomato-positive neurons in these areas were co-labeled with c-Fos (fig. S6, D and E-I). Together, our data suggest that sTRN receives inputs from wide brain regions that are associated with stress information processing.

## Discussion

In this study, we report a previously unknown function of the TRN in the thalamic control of depression. We reported three major findings. First, the LHb^GLU^ neurons receive inhibitory input from sensory TRN which can be activated by acute aversive stimuli. Second, chronic restraint stress attenuates the synaptic strength of the sTRN^SOM^-LHb circuit, which underlies the hyperactivity of LHb neurons and depression onset. Last, the sTRN^SOM^-LHb pathway specifically restores depressive-like behaviors induced by chronic stress and chronic pain, rather than pain-like behaviors induced by nerve injury and depression.

### sTRN-LHb monosynaptic inhibitory projections

Thalamic inhibition is a critical element of thalamic projecting neuron modulation, and its perturbation is found in many diseases (*28*). The TRN is one of the major sources of thalamic inhibition (*26*). The neurons within TRN are highly heterogeneous in anatomical distribution, molecular identities, electrophysiological properties, synaptic connectivity, and function (*35*, *38*-*41*). Although previous studies have reported hierarchical connectivity between TRN and distinct thalamic areas, for example, the core region of TRN projects to first-order thalamic nuclei whereas the shell region of TRN projects to the high-order thalamic nuclei (*35*), and dorsal TRN predominantly projected to the posterior thalamic nucleus (Po) whereas the ventral TRN mainly innervated the ventrobasal thalamus (VB) (*33*), the anatomical connectivity and function of TRN-LHb projections are not yet reported.

We performed cell-specific anterograde tracing experiments and observed that PV^+^ and SOM^+^ neurons in the sTRN have different projection areas. SOM^+^ neurons of the caudal sector of TRN (referred to as sTRN) broadly projected to LHb, ventrolateral part of the laterodorsal thalamus (LDVL), mediorostral part of the lateral posterior thalamus (LPMR), VB, Po and ventrolateral thalamus (VL). Further retrograde monosynaptic tracing and electrophysiological results confirmed the inhibitory projections of sTRN^SOM^-LHb. In contrast, PV^+^ neurons of the sTRN predominantly projected to VB, Po, and VL, with sparse projection to LDVL and LPMR, and without projection to LHb. Interestingly, the innervation distribution by two types of neurons displayed somewhat non-overlapping, even for VB, Po, and VL. This complementary innervation of the dorsal thalamus by molecularly diverse neurons within sTRN enriches the understanding of a comprehensive architecture of the TRN-thalamic nuclei connectivity and could be the neural substrates for segregated emotion (i.e., depression) and somatosensation (i.e., pain) modulation. The suspect is supported by our data showing that TRN^SOM^-LHb projections modulate depression rather than pain, and by previous studies showing that the TRN-VB projections contribute to pain regulation (*33*, *42*). It remains to be determined whether the TRN-VB circuit also regulates depression and the cellular identity of TRN-VB projections that modulate pain.

### sTRN^SOM^-LHb projections in depression onset under chronic stress

Despite much evidence on the involvement of LHb aberrant hyperactivity in depressive disorder (*9*-*11*, *13*), the neural circuit mechanisms underlying abnormal LHb activity and depression under stress have remained elusive, especially from the inhibitory circuit perspective. By using *in vivo* calcium imaging, we demonstrated that LHb-projecting inhibitory sTRN afferents are activated by aversive stimuli including acute physical restraint suggesting their involvement in the aversive emotion encoding which has not been reported previously. We speculated that this adaptive response possibly helps to protect organisms against acute stress. However, maladaptive hypoexcitability of TRN-LHb projections occurs during chronic stress. *In vitro* electrophysiological recordings revealed that CRS blunts the synaptic strength of the sTRN^SOM^-LHb circuit. To further examine the TRN-LHb inhibitory projections in the pathophysiology of depression, we repetitively activate LHb-upstream sTRN neurons during CRS, and intriguingly found that both CRS-induced LHb neural hyperactivity and depression onset were prevented. Moreover, artificially silencing the sTRN^SOM^-LHb circuit is sufficient to induce depressive-like behaviors in naïve mice. These data collectively suggest that the attenuation of TRN-LHb projections during CRS underlies depression onset. In combination with previously well-elucidated enhanced excitatory inputs to LHb during chronic stress, (*9*, *19*, *20*), our findings advance a better understanding of excitation/inhibition imbalance in hyperactive LHb neural activity and pathophysiology of depression.

### sTRN^SOM^-LHb projections restore depression in chronic stress and chronic pain

Here, we tested two independent depressive-like mouse models: chronic restraint stress-induced depression and chronic pain-induced comorbid depression. Either specific activation of LHb-projecting sTRN^SOM^ afferents or inhibition of sTRN-targeted postsynaptic LHb neurons is sufficient to rescue the depressive-like behaviors in the above two models. Given the involvement of LHb in processing pain information and analgesic signals (*15*-*17*), we also assessed the role of the sTRN-LHb pathway in pain regulation. Unexpectedly, the same manipulation of the sTRN-LHb circuit does not affect pain in both the nerve injury-induced pain model and the CRS-induced comorbid pain model. The segregation control of depression and pain might arise from the possibility that depression-specific ensembles in the LHb, rather than pain-specific neurons, are targeted by sTRN. Future activity-dependent cellular labeling systems should be warranted to dissect the different subpopulations of LHb neurons processing pain and depression, and their respective connections with TRN.

In addition, previous studies reported that PV^+^ and SOM^+^ neurons in TRN receive projections from brain areas that are mainly related to sensory and emotional processing, respectively (*38*, *41*). Furthermore, the divergent control of different events could also arise from distinct synaptic inputs and intrinsic physiological characteristics of TRN cell types, shaping distinct spiking outputs and thus tuning to discrete behavioral events (*38*, *41*). We dissected upstream brain regions of sTRN which were activated by stress, including CeA/BLA, ACC, IC, and DRN. These areas were all implicated in emotion encoding (*43*-*48*). Where is the upstream brain areas of sTRN associated with pain and how cell type-based circuit-specific temporal tuning is involved in depression and pain processing, are interesting.

Overall, our study revealed the dysfunctional adaptation of a novel GABAergic sTRN**-**LHb pathway during chronic stress which significantly bridges the gap in the clarification of inhibitory circuit mechanisms underlying the pathophysiology of depression. We also demonstrated that repetitive sTRN-LHb circuit activation during chronic stress can prevent depression onset, and transient sTRN-LHb circuit activation can alleviate depression in chronic stress or chronic pain, shedding light on early intervention targets before depression onset and therapeutic targets for depression management.

## Acknowledgments

This study was supported by the National Natural Science Foundation of China (grants 32271048 and 32070999 to Yan Z, U20A20357 to X. F. L, 31972905 and 32271190 to Ying Z.) and Anhui Provincial Natural Science Foundation (grant 2008085J16 to Yan Z.).

## Author contributions

X. X., R. C., and X. Y. W. designed the studies and conducted most of the experiments and data analysis. X. X. and R. C. wrote the first draft. W. B. J. and P. F. X generated some behavioral data. X. Q. L. managed the mouse colonies used in this study. Ying Z., X. F. L. and Yan Z. were involved in the overall design of the project and editing of the final manuscript.

## Declaration of interests

The authors declare no competing interests.

All data necessary to understand and assess the conclusions of this study are available in the main text or the supplementary materials. There are no restrictions on data availability in the manuscript.

## Methods

### Animals

Adult (8-10 weeks) C57BL/6J, *SOM-Cre*, *PV-Cre*, *CaMKIIα-Cre*, and *Rosa*26-tdTomato *(Ai14)* male mice were used in this study. *C57BL/6J* mice were purchased from Beijing Vital River Laboratory Animal Technology, and *Ai14*, *SOM-Cre*, *PV-Cre*, and *CaMKIIα-Cre* mice were initially acquired from the Jackson Laboratory. Animals were maintained at a stable room temperature (23 ± 1 °C) with a 12-hour light/dark cycle (lights on from 7:00 to 19:00). They were housed five per cage in a colony with ad libitum water and food. All animal experiment procedures were approved by the animal care and use committee of the University of Science and Technology of China.

### Animal models

#### Chronic restraint stress model (CRS)

Mice were placed in restraint tubes made of 50 ml centrifuge tubes for 6 hours (from 9:00 to 15:00) per day for consecutive 21 days. The centrifugal tube is pre-stamped with air holes scattered throughout the tube to allow the mouse to breathe, and there is a small hole in the center of the lid through which the mouse’s tail is exposed to air (see Fig. S3A). During the restraint period, control mice were allowed to freely move around the cage, but fasted for water and food. At the end of 21 days, the mice were allowed to take 1 day off to exclude the effects of acute stress.

#### Spared nerve injury model (SNI)

Spared nerve was injured following the protocol previously described (*49*). In brief, the mice were anesthetized with 3% isoflurane. An incision was made in the middle of the left thigh using the femur as a marker. The skin and muscle were incised to explore the sciatic nerve comprising the sural, common peroneal, and tibial nerves. The tibial and common peroneal nerves were ligated with nonabsorbent 4-0 chromic gut and transected (1-2 mm slices), while the sural nerve was carefully preserved. The skin was sutured with 6-0 silk and disinfected by iodophor. For the sham group, the procedure was the same as for the experimental group except for ligation and transection of the nerves.

#### Stereotaxic surgeries

Before surgery, the mice were fixed in a stereotactic frame (RWD, Shenzhen, China) under anesthesia with 3% pentobarbital sodium (30 mg/kg) intraperitoneally (i.p.), and body temperature was maintained with a heating pad. Erythromycin eye ointment was applied to maintain eye lubrication. The skull surface was exposed with a midline scalp incision, and the injection site was defined using stereotactic coordinates (see below). An adental drill (RWD, DC 30V) was used for craniotomy (∼0.5 mm hole) to enable virus injection. Injections of 150-200 nl virus (depending on the expression strength and viral titer) were carried out through the glass pipettes (tip diameter of 10-30 μm) connected to an infusion pump (KD Scientific) at a rate of 50 nl/min. After the injection is completed, the glass pipettes were left for 10 min before withdrawal to allow virus diffusion. The mice were allowed to recover from anesthesia on a heating blanket before returning to the home cage. The coordinates were defined as dorso-ventral (DV) from the brain surface, anterior-posterior (AP) from bregma and medio-lateral (ML) from the midline (in mm).

#### Virus injection and optical fibers implantation

For pharmacogenetic inhibition or activation of LHb glutamatergic neurons, C57BL/6J mice were bilaterally microinjected with 150 nl of rAAV2/9-CaMKIIα-hM4Di-mCherry-WPREs (titer: 5.91E+12 vg/ml, BrainVTA, PT-0050) or rAAV2/9-CaMKIIα-hM3Dq-mCherry-WPREs (titer: 5.29E+12 vg/ml, BrainVTA, PT-0049), and rAAV2/9-CaMKIIα-mCherry-WPRE-pA (titer: 5.14E+12 vg/ml, BrainVTA, PT-0108) as a control into the LHb. For pharmacogenetic inhibition or activation of sTRN-targeted LHb neurons, C57BL/6J mice were bilaterally microinjected with 150 nl of rAAV2/1-hSyn-Cre-WPRE-pA (titer: 1.00E+12 vg/mL, BrainVTA, PT-0136) into sTRN, and 150 nl of rAAV2/9-EF1α-DIO-hM4Di-mCherry-WPREs (titer: 5.18E+12 vg/ml, BrainVTA, PT-0043) or rAAV2/9-Ef1α-DIO-hM3Dq-mCherry-WPREs (titer: 5.27E+12 vg/ml, BrainVTA, PT-0042). For pharmacogenetic activation of LHb-projecting sTRN neurons, C57BL/6J mice were bilaterally microinjected with 150 nl of rAAV2/2-retro-hSyn-Cre-WPRE-pA (titer: 5.22E+12 vg/ml, BrainVTA, PT-0136) into the LHb and the rAAV2/9-Ef1α-DIO-hM3Dq-mCherry-WPREs (titer: 5.27E+12 vg/ml, BrainVTA, PT-0042) was subsequently delivered into the sTRN. rAAV2/9-EF1α-DIO-mCherry-WPRE-pA (titer: 5.14E+12 vg/ml, BrainVTA, PT-0013) was used as a control. For chemogenetic manipulation of neuronal activity, mice were injected with clozapine N-oxide (CNO, i.p., 2 mg/kg, APExBIO Technology LLC).

For optogenetic stimulation, mice were bilaterally microinjected with 150 nl of rAAV2/9-EF1α-DIO-hChR2(H134R)-mCherry-WPRE-pA (titer: 4.50E+12 vg/ml, BrainVTA, PT-0002) or rAAV2/9-EF1α-DIO-eNpHR3.0-EYFP-WPRE-pA (titer: 5.36E+12 vg/mL, BrainVTA, PT-0006) into the sTRN.

For monosynaptic anterograde tracing, we microinjected 150-250 nl of rAAV2/1-hSyn-Cre-WPRE-pA (titer: 1.00E+12 vg/mL, BrainVTA, PT-0136) into the unilateral sTRN of C57BL/6J mice. The rAAV2/9-EF1α-DIO-mCherry-WPRE-pA (titer: 5.14E+12 vg/mL, BrainVTA, PT-0013) was subsequently delivered into the LHb. For monosynaptic retrograde tracing, we microinjected 150-200 nl of rAAV2/2-retro-hSyn-Cre-WPRE-pA (titer: 5.22E+12 vg/ml, BrainVTA, PT-0136) into the unilateral LHb of C57BL/6J mice. The rAAV2/9-EF1α-DIO-mCherry-WPRE-pA (titer: 5.14E+12 vg/ml, BrainVTA, PT-0013) was subsequently delivered into the sTRN. For sTRN upstream brain region dissection, 150 nl of rAAV2/2-retro-hSyn-Cre-WPRE-pA (titer: 5.77E+12 vg/ml, BrainVTA, PT-0136) was injected into sTRN of *Ai14* mice.

#### Optical fibers implantation

For optogenetic manipulation in awake behaving mice, chronically optical fiber (diameter: 200 μm; N.A., 0.37; length, 4 mm; Inper) was implanted 200 (m above the virus injection site in the bilateral LHb (AP: -1.9 mm; ML: ± 1.1mm; DV: -2.45 mm at an angle of 15°). The fibers were attached to the skull with dental cement and connected to a laser generator using an optical fiber sleeve. The delivery of blue light (473 nm, 1-3 mW, 15 ms pulses, 20 Hz) or yellow light (594 nm, 5-8 mW, 15 ms pulses, 20 Hz) was controlled by a class laser product (QAXK-LASER, ThinkerTech). The same stimulus protocol was applied in the control group. Following surgery, the mice were allowed to recover for at least 1 week before performing the behavioral experiments. We checked the location of fibers after all experiments and discarded the data obtained from mice in which the fibers were outside the desired brain region.

#### Fiber photometry recording

The three-color single-channel fiber photometry system (ThinkerTech) was used for recording Ca^2+^ signals from LHb neurons or LHb-projecting TRN afferents. The 150 nl of rAAV2/9-hSyn-GCaMP6m-WPRE-pA (titer: 5.51E+12 vg/ml, BrainVTA, PT-0148) virus was injected into the sTRN (AP: -1.5 mm; ML: +2.35 mm; DV: -3.25 mm) of C57BL/6J mice. The 150 nl of rAAV2/9-DIO-GCaMP6m-WPRE-pA (titer: 6.21E+12 vg/ml, BrainVTA, PT-0283) virus was injected into the LHb (AP: -1.9 mm; ML: +0.45 mm; DV: -2.55 mm) of *CaMKIIα-Cre* mice. An optical fiber (diameter: 200 μm; N.A., 0.37; length, 4 mm; Inper) was subsequently implanted into the sTRN or LHb. The optical fiber was affixed with a skull-penetrating screw and with dental acrylic. To enable recovery and AAV expression, mice were housed individually for at least 10 days following virus injection. After three weeks, the fiber photometry data were recorded continuously during the air puff, pinch and physical restraint tests. The normalized Delta F/F values and traces were visualized using custom MATLAB (MathWorks) scripts that were produced by ThinkerTech. In addition, we excluded the mice that had no GCaMP6m signals in response to air puff stimuli before performing tests, which may be due to the failure of viral expression and missed targets, including the injection of viruses, placement of optical fiber or optical fiber tip clogging.

#### Depression-related behaviors test

For all behavioral tests, dim light (20 lux) and a quiet environment were used in the room to minimize the anxiety of the animals. The behavioral experiments described herein were performed by experimenters who were blind to the treatments.

#### Open field test (OFT)

Motor activity was tested in open field test box (40×40×40cm). Individual mice were introduced into the center of the box in a room with dim light and were allowed to freely explore their surroundings during 6 min test session with a video-tracking system. The total distance traveled in the last 5 min was analyzed by SMART V3.0 software (Panlab S.L., Spain). To remove olfactory interference, we cleaned the box with 75% ethanol after each test. To examine the effect of the optogenetic manipulation on locomotion, the total distance traveled in the first 10 min (with the 5 min light-off period followed by the 5 min light-on period) was analyzed.

#### Forced swim test (FST)

Mice were individually placed into a transparent plexiglass cylinder (diameter 12 cm, height 30 cm), containing a height 20 cm of water at 24 ± 2℃. In the test, the time of swimming and immobility was recorded during a 6 min period. The processes were videotaped from the side. Immobility was assigned when no additional activity was observed other than that required to keep the mice head above the water. The time that mice spent in immobility in the last 4 min was quantified offline manually by an observer blinded to animal treatment. Animals were never allowed to drown during the test. Mice with nerve injury were exempted from the FST because of the undermined swimming skill.

#### Tail suspension test (TST)

Mice were suspended by the tip of tail using adhesive tape. One end of the tape was attached to a horizontal table that surface above 30 cm from the floor. During the 6 min process, the behavior was videotaped from the side. The immobile time during the last 5 min was manually recorded by an observer blinded to animal treatment. Mice were considered immobile when they were completely motionless or passive swaying. For optogenetic manipulations, mice were first hung by the tail for 2 min, followed by a 3 min off/ 3 min on/ 3 min off light epoch.

#### Sucrose preference test (SPT)

The animals were housed in a single cage and given two bottles of distilled water for 48 h training, then two bottles of 1% sucrose water for 48 h training. After 24h of water deprivation, the final test was performed for 2 h, giving one bottle of water and one bottle of 1% sucrose water. The positions of the two water bottles were exchanged halfway through the period of 1 h. The sugar-water preference index was calculated as the ratio of 1% sucrose water to the total water (water and sucrose water) consumed. For the optogenetic experiment, mice were subjected to a 30-minute light period and sugar-water preference index was measured.

#### Mechanical allodynia assay

Before behavioral testing, all animals were habituated to plexiglass chambers (6.5×6.5×6 cm) positioned on a wire mesh grid for at least two days. Once the mice were calm, the plantar area of hind paws was stimulated with a series of Von Frey filaments with different strengths (g) to measure the mechanical withdrawal threshold. The stimulus producing a 50% likelihood of a withdrawal response was determined and taken as the paw withdrawal threshold (PWT) using the Up-Down method (*50*).

#### Immunofluorescence and imaging

Mice were deeply anesthetized with 3% isoflurane and then intracardially perfused with 20 ml 0.01M phosphate-buffered saline (PBS) (4℃) and 20 ml 4% paraformaldehyde (PFA) in PBS (4℃). After post-fixation overnight, the mouse brain was cryoprotected with 30% sucrose for two days. The brain was embedded in OCT (SAKURA, 4583) and sectioned coronally (30 μm thick) with a freezing microtome (Leica CM1950). For immunofluorescent staining, the sections were blocked with 3% BSA in PBS with 0.3% Triton X-100 for 1h at room temperature and subsequently incubated with primary antibodies (rabbit anti-glutamate, 1:500, Sigma; rabbit anti-SOM, 1:500, Novus; rabbit anti-c-Fos, 1:500, Synaptic Systems) at 4°C overnight. After washing with PBS, the sections were subsequently coupled with the corresponding fluorophore-conjugated secondary antibodies (1:500, Jackson) for 1.5 h at room temperature. Finally, after washing 3 times for 7 min in PBS, sections were stained with DAPI, and the slides were sealed with anti-fluorescence quenching sealing tablets. Images were captured with Olympus confocal microscopes (FV3000, Olympus) and analyzed with ImageJ software.

#### Brain slice electrophysiology Brain slice preparation

For slices preparation, mice were anesthetized with isoflurane followed by pentobarbital (30 mg/kg, i.p.) and intracardially perfused with ∼20 ml oxygenated ice-cold modified N-methyl-D-glucamine and artificial cerebrospinal fluid (NMDG ACSF) that contained (in mM) 93 NMDG, 2.5 KCl, 1.2 NaH_2_PO_4_, 30 NaHCO_3_, 20 N-2-hydroxyethylpiperazine-N-2-ethanesulfonic acid (HEPES), 25 glucose, 5 Na-ascorbate, 3 Na-pyruvate, 0.5 CaCl_2_, 10 MgSO_4_, 3 glutathione (GSH) and 2 thiourea (pH: 7.3-7.4, osmolarity: 300-310 mOsm). Coronal slices (300 µm) containing the sTRN or LHb were sectioned in chilled (2-4℃) NMDG ACSF at 0.18 mm/s velocity on a vibrating microtome (VT1200s, Leica). The brain slices were initially incubated in NMDG ACSF (saturated with 95% O_2_/5% CO_2_ to provide a stable potential of hydrogen and continuous oxygenation) for 10 min at 33 °C, followed by (HEPES) ACSF that contained (in mM) 92 NaCl, 2.5 KCl, 1.2 NaH_2_PO_4_, 30 NaHCO_3_, 20 HEPES, 25 glucose, 2 thiourea, 5 Na-ascorbate, 3 Na-pyruvate, 2 CaCl_2_, 2 MgSO_4_ and 3 GSH (pH 7.3-7.4, osmolarity 300-310 mOsm) for at least 1 h at 25 °C. The brain slices were transferred to a slice chamber for electrophysiological recording and were continuously submerged and superfused with standard ACSF that contained (in mM) 129 NaCl, 2.4 CaCl_2_, 3 KCl, 1.3 MgSO_4_, 20 NaHCO_3_, 1.2 KH_2_PO_4_ and 10 glucose (pH 7.3-7.4, osmolarity 300-310 mOsm) at 5 ml/min at room temperature. During recording and analysis, the recorders were blind to group identity.

#### Electrophysiological verification of pharmacogenetics and optogenetics

Whole-cell patch-clamp recordings were obtained from visually identified sTRN or LHb cells. Neurons in the slice were visualized using a ×40 water-immersion objective on an upright microscope (BX51WI, Olympus) equipped with infrared-differential interference contrast (IR-DIC) and an infrared camera connected to the video monitor. Patch pipettes (3-5 MΩ) were pulled from a horizontal puller (P1000, Sutter Instruments) and filled with the internal solution that contained (in mM): 130 K-gluconate, 5 KCl, 4 Na_2_ATP, 0.5 NaGTP, 20 HEPES, 0.5 EGTA, (PH 7.28, 290-300 mOsm). Whole-cell patch-clamp recordings were acquired via a Multiclamp 700B patch-clamp amplifier (Molecular Devices) and Digidata 1550B (Molecular Devices), digitized at 5 kHz and filtered at 2 kHz. Data were analyzed with pClamp 10.0 software. Cells were excluded when series resistance changed more than 20% during the recording.

To confirm the efficacy of ChR2-mediated activation, fluorescently labeled neurons that expressed ChR2 in *SOM-Cre* mice 3 weeks after virus injection were visualized and stimulated with a 473 nm laser (QAXK-LASER, China) using 20 Hz stimulation protocols with a pulse width of 15 ms. After a stable membrane potential was acquired, 473-nm laser illumination induced reliable spikes in TRN neurons. Similarly, the functional expression of eNpHR3.0 was assessed by applying yellow (594 nm) laser light stimulation. To confirm the efficacy of hM4Di-mediated inhibition and hM3Dq-mediated excitation, CNO (10 μM, APExBIO Technology LLC) was bath-applied.

To examine the firing rate of LHb neurons, 500 ms pulses with 10 pA command current steps were injected from -60 to +150 pA, and the numbers of spikes were quantified for each step.

To examine functional inhibitory projections from the sTRN to the LHb, membrane potentials of LHb neurons were held at 0 mV to record light-evoked IPSCs. For evaluating synaptic identities, GABA-mediated IPSCs were blocked by the bath application of bicuculline (10 μM, Sigma). To test direct synaptic connections, both TTX (1 μM, Sigma) and 4-AP (500 μM, Sigma) were used to restore monosynaptic current. For evaluating the presynaptic mechanism, paired pulses (15 ms duration) with an interval of 50 ms (ISI 50 ms) were delivered, and the PPR was calculated as the amplitude ratio IPSC_2_/IPSC_1_.

To examine miniature IPSCs (mIPSCs) of LHb neurons, patch pipettes were filled with the chloride-based internal solution that contained (in mM): 145 CsCl, 10 EGTA, 10 HEPES, 2 MgCl2, 2 CaCl2, 2 Mg-ATP, and the membrane potentials of LHb neurons were held at -70 mV, resulting in inward mIPSCs. Moreover, TTX (1 μM) and CNQX (10 μM, Sigma) were added to eliminate spontaneous action potentials and AMPAR-mediated inward mEPSCs, respectively.

#### Quantification and statistical analysis

All experiments and data analyses were conducted blindly, including the immunohistochemistry, electrophysiology and behavioral analyses. Data were analyzed with GraphPad Prism v.8.0.1, Olympus FV10-ASW 4.0a Viewer, Microsoft office 2021, and MATLAB R2016a software. Normality was assessed using the Shapiro-Wilk test. When normally distributed, the data were analyzed with paired t-tests, unpaired t-tests as appropriate. When normality was violated, the data were analyzed with Wilcoxon signed-rank test for paired test and Mann-Whitney U test for unpaired test. Behavioral data were analyzed by one-way or two-way analysis of variance (ANOVA) followed by Bonferroni’s test for multiple comparisons, and the unpaired Student’s t-test or Mann-Whitney U test for two group comparisons. Imaging of calcium activity data were analyzed by paired Student’s t-test or Wilcoxon signed-rank test. For electrophysiological results, data were assessed by two-way analysis of variance (ANOVA) followed by Bonferroni’s test for multiple comparisons, and the unpaired Student’s t-test or Mann-Whitney U test for two group comparisons. For immunofluorescence analysis, data were analyzed using unpaired Student’s t-test or Mann-Whitney U test. Statistical significances were represented as *p < 0.05, **p < 0.01, ***p < 0.001, ****p < 0.0001; ^#^p < 0.05, ^##^p < 0.01, ^###^p < 0.001, ^####^p < 0.001. All data were expressed as mean ± standard error of means (S.E.M.) except for data in Figure 1E and 1G shown as box and whisker plots.

## Notes

### Competing Interest Statement

The authors have declared no competing interest.

### Summary of Updates

Added a new set of data showing chronic restraint stress (CRS) weakens sTRN-LHb synaptic strength as new Figure 2, and revised the text part correspondingly.

