## Supplemental figures for "The thalamic reticular nucleus-lateral habenula circuit regulates depressive-like behaviors in chronic stress and chronic pain"

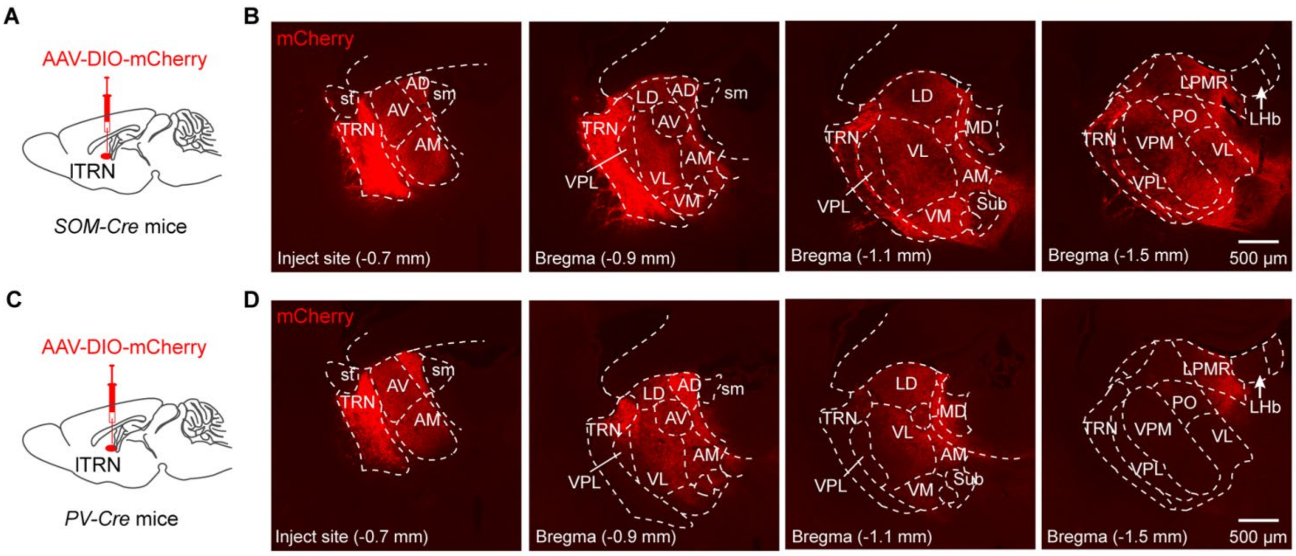


**Fig. S1. Cre-dependent anterograde tracing of limbic TRN neurons. Related to Fig. 1.**

(A and C) Schematic showing viral injection into the limbic TRN of *SOM-Cre* (A) or *PV-Cre* (C) mice.

(B and D) Representative images of mCherry-expressing signals in different brain regions of *SOM-Cre* (B) or *PV-Cre* (D) mice. Scale bars, 500 μm.

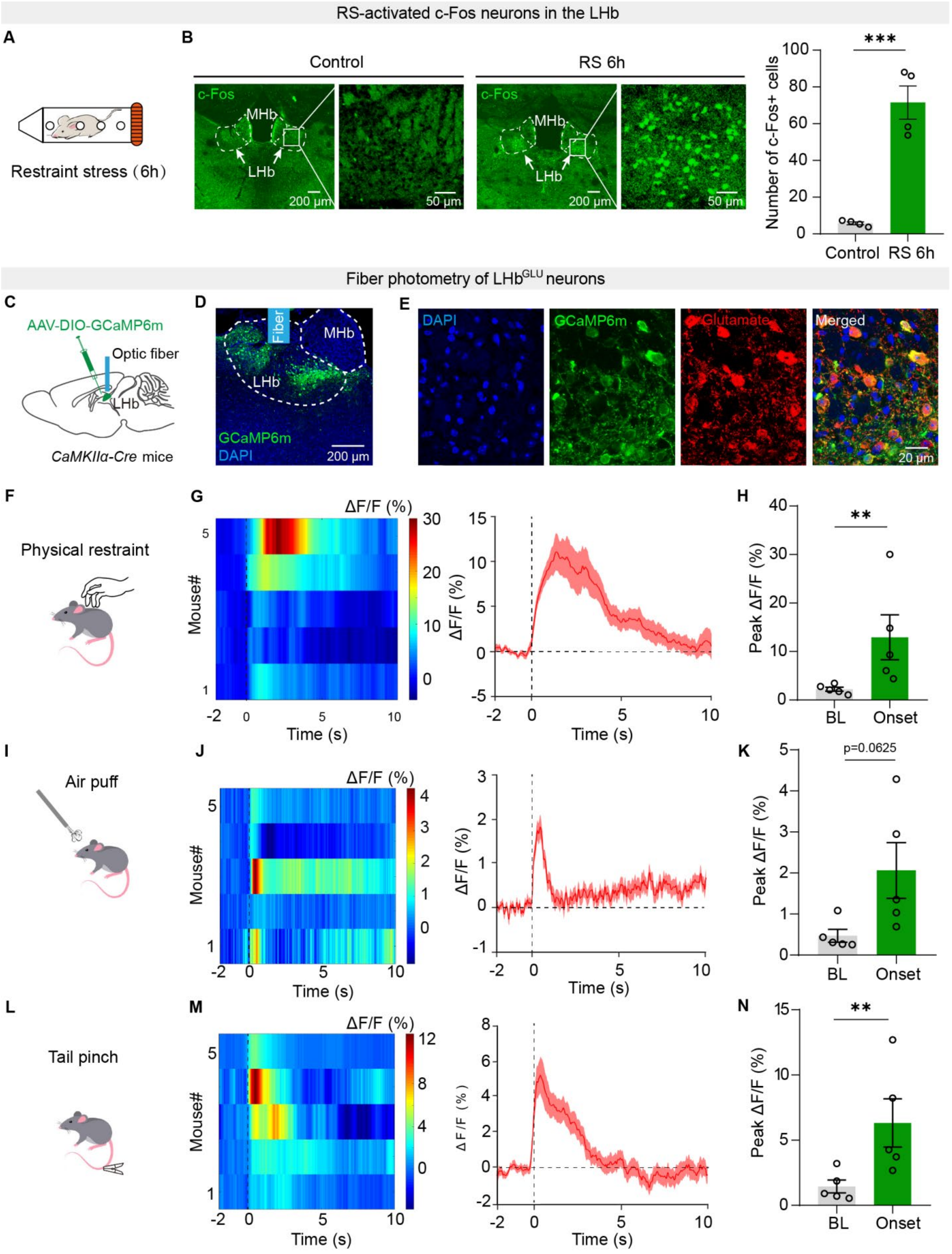


**Fig. S2. Aversive stimuli activate LHb$^{GLU}$ neurons. Related to Fig. 1.**

(A) Schematic of restraint stress (RS).

(B) Representative images (left) and quantification (right) of c-Fos staining in

RS-treated or control mice. Scale bars, 200 μm and 50 μm. (n=4 mice/group)

(C and D) Illustration (C) and representative image (D) of viral delivery and optic fiber implantation. Scale bar, 200 μm.

(E) Representative images validating specific expression of GCaMP6m in $LHb^{GLU}$ neurons. Scale bar, 20 μm.

(F, I and L) Schematic of fiber photometry recording in response to RS (F), air puff (I), and tail pinch (L).

(G, J and M) Heatmap and average responses showing $Ca^{2+}$ transients evoked by RS (G), air puff (J), and tail pinch (M) in the LHb neurons. (n=5 mice/group)

(H, K and N) Quantification of peak average $Ca^{2+}$ responses before and after RS (H), air puff (K), and tail pinch (N) stimulation. (n=5 mice/group)

Data are presented as mean ± SEM. **$p < 0.01$, ***$p < 0.001$. Unpaired two-sided t test for (B); Wilcoxon signed-rank test for (K); Paired ratio two-sided t test for (H) and (N).

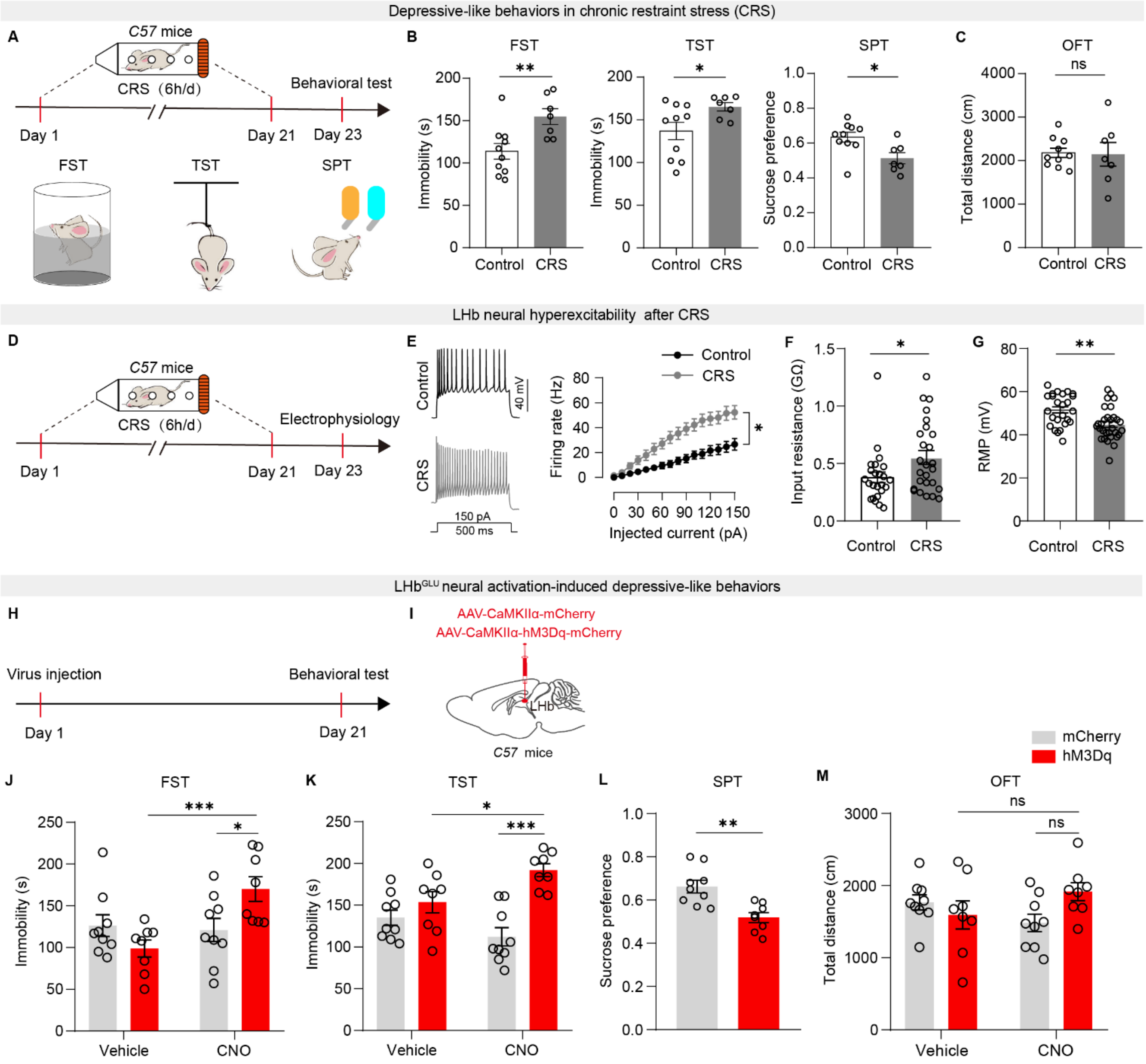


**Fig. S3. CRS induces LHb neural hyperexcitability and LHb neural activation induces depressive-like behaviors. Related to Fig. 2, Fig. 3 and Fig. 4.**

(A, D and H) Schematic of experimental design.

(B and C) Depressive-like behaviors assessed by FST, TST, and SPT (B), and locomotion activity assessed by OFT (C) in control and CRS-treated *C57* mice. (n=7-10 mice/group)

(E-G) Summary data for the firing rate (E), input resistance (F), and resting membrane potential (G) recorded from LHb neurons of control or CRS-treated *C57* mice.

(I) Illustration of viral delivery into the LHb.

(J-M) Behavioral effects of chemogenetic activation of LHb neurons on depressive-like behaviors assessed by FST (J), TST (K), and SPT (L), and locomotion assessed by OFT (M) in naïve *C57* mice. (n=8-9 mice/group)

Data are presented as mean ± SEM. “ns”, no significance; *$p < 0.05$, **$p < 0.01$, ***$p < 0.001$. Unpaired two-sided t test for (B), (C), (G), and (L); Two-way repeated measures ANOVA with Bonferroni *post hoc* analysis for (E), (J), (K), and (M); Mann-Whitney U test for (F).

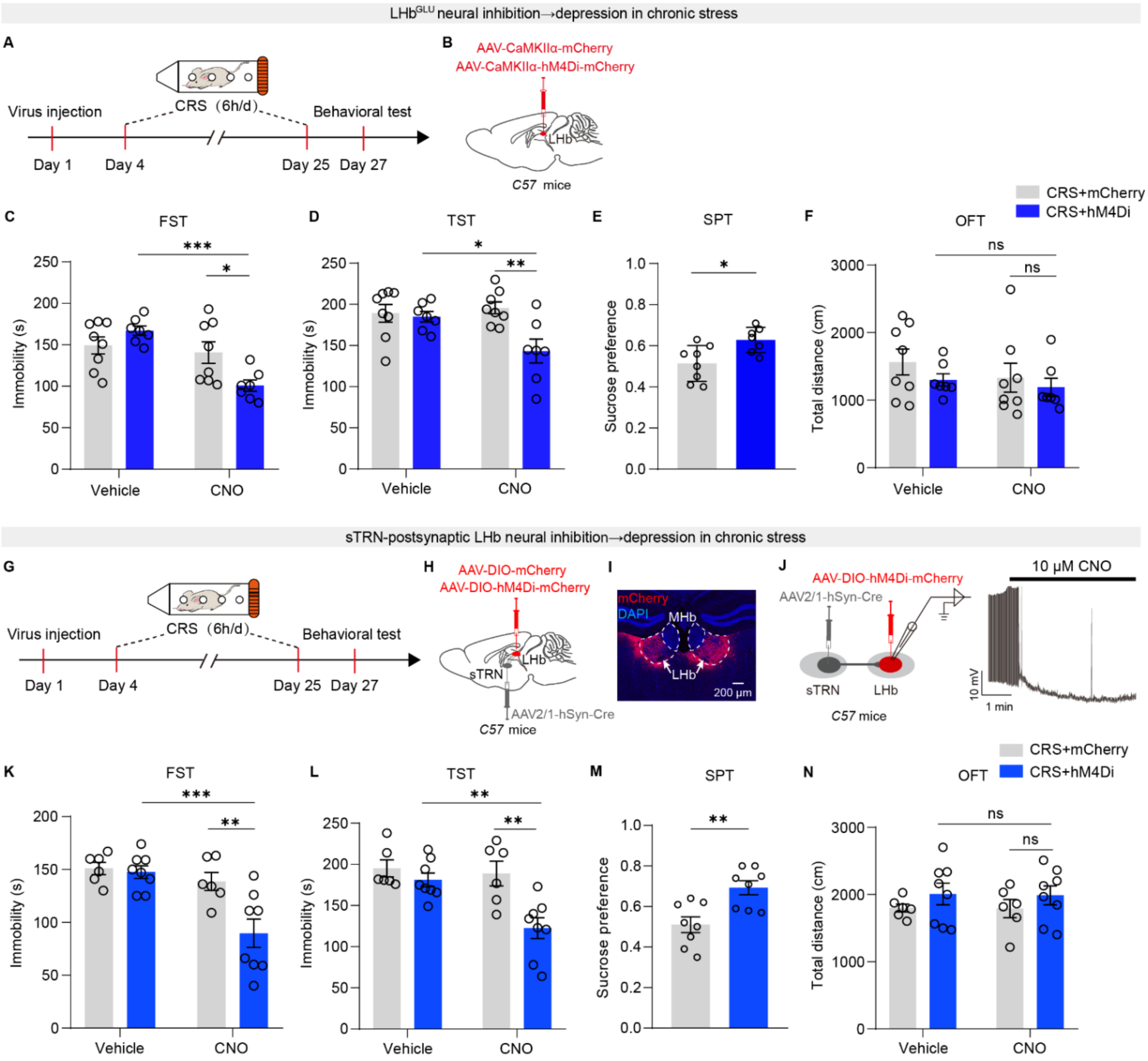


**Fig. S4. Chemogenetic inhibition of sTRN-postsynaptic LHb neurons alleviates depressive-like behaviors induced by chronic stress. Related to Fig. 5.**

(A and G) Schematic of experimental design.

(B) Illustration of viral delivery into the LHb.

(C-F) Behavioral effects of chemogenetic inhibition of LHb neurons on depressive-like behaviors assessed by FST (C), TST (D), and SPT (E), and locomotion assessed by OFT (F) in naïve *C57* mice. (n=7-8 mice/group)

(H and I) Illustration (H) and representative image (I) of viral delivery. Scale bar, 200 µm.

(J) Schematic of recording configuration in acute slices and representative trace showing hypolarization of membrane potential in a hM4Di-mCherry$^+$ LHb neuron by CNO (10 μM).

(K-N) Behavioral effects of chemogenetic inhibition of LHb neurons on depressive-like behaviors assessed by FST (K), TST (L), and SPT (M), and locomotion activity assessed by OFT (N) in CRS-treated *C57* mice. (n=6-8 mice/group)

Data are presented as mean ± SEM. “ns”, no significance; *p < 0.05, **p < 0.01, ***p < 0.001. Unpaired two-sided t test for (E) and (M); Two-way repeated measures ANOVA with Bonferroni *post hoc* analysis for (C)-(F) and (K)-(N).

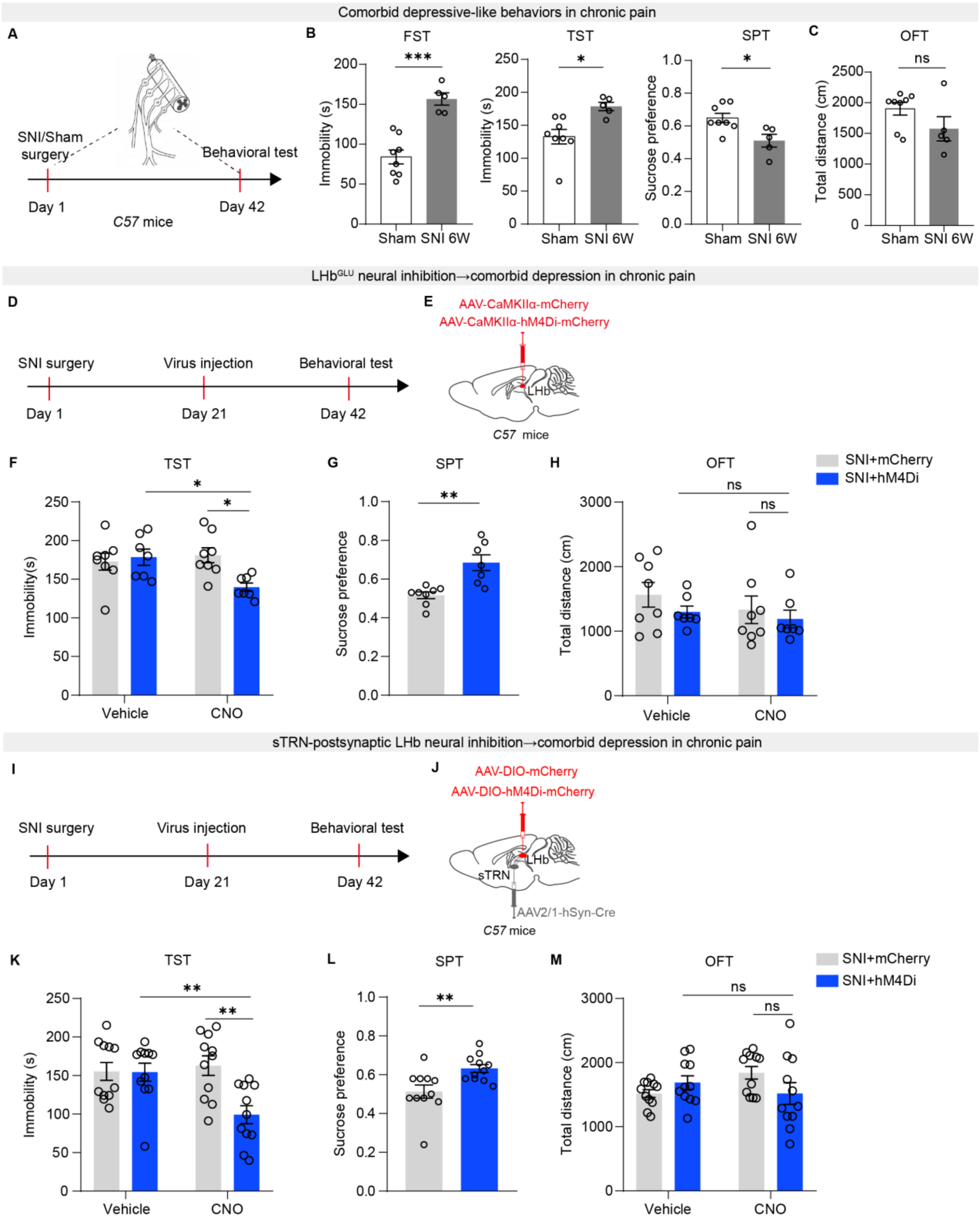


69 **Fig. S5. Chemogenetic inhibition of sTRN-postsynaptic LHb neurons alleviates**

70 **depressive-like behaviors induced by chronic pain. Related to Fig. 5.**

71 (A, D and I) Schematic of experimental design.

72 (B and C) Depressive-like behaviors assessed by FST, TST, and SPT (B), and

73 locomotion activity assessed by OFT (C) in sham and SNI-treated *C57* mice. (n=5-8

mice for each group)

(E and J) Illustration of viral delivery.

(F-H) Behavioral effects of chemogenetic inhibition of LHb neurons on depressive-like behaviors assessed by TST (F) and SPT (G), and locomotion activity assessed by OFT (H) in SNI-treated *C57* mice. (n=7-8 mice/group)

(K-M) Behavioral effects of chemogenetic inhibition of LHb neurons on depressive-like behaviors assessed by TST (K) and SPT (L), and locomotion activity assessed by OFT (M) in SNI-treated *C57* mice. (n=11 mice/group).

Data are presented as mean ± SEM. “ns”, no significance; *p < 0.05, **p < 0.01, ***p < 0.001. Two-way repeated measures ANOVA with Bonferroni *post hoc* analysis for (F), (H), (K), and (M); Unpaired two-sided t test for (B) and (G); Mann-Whitney U test for (C) and (L).

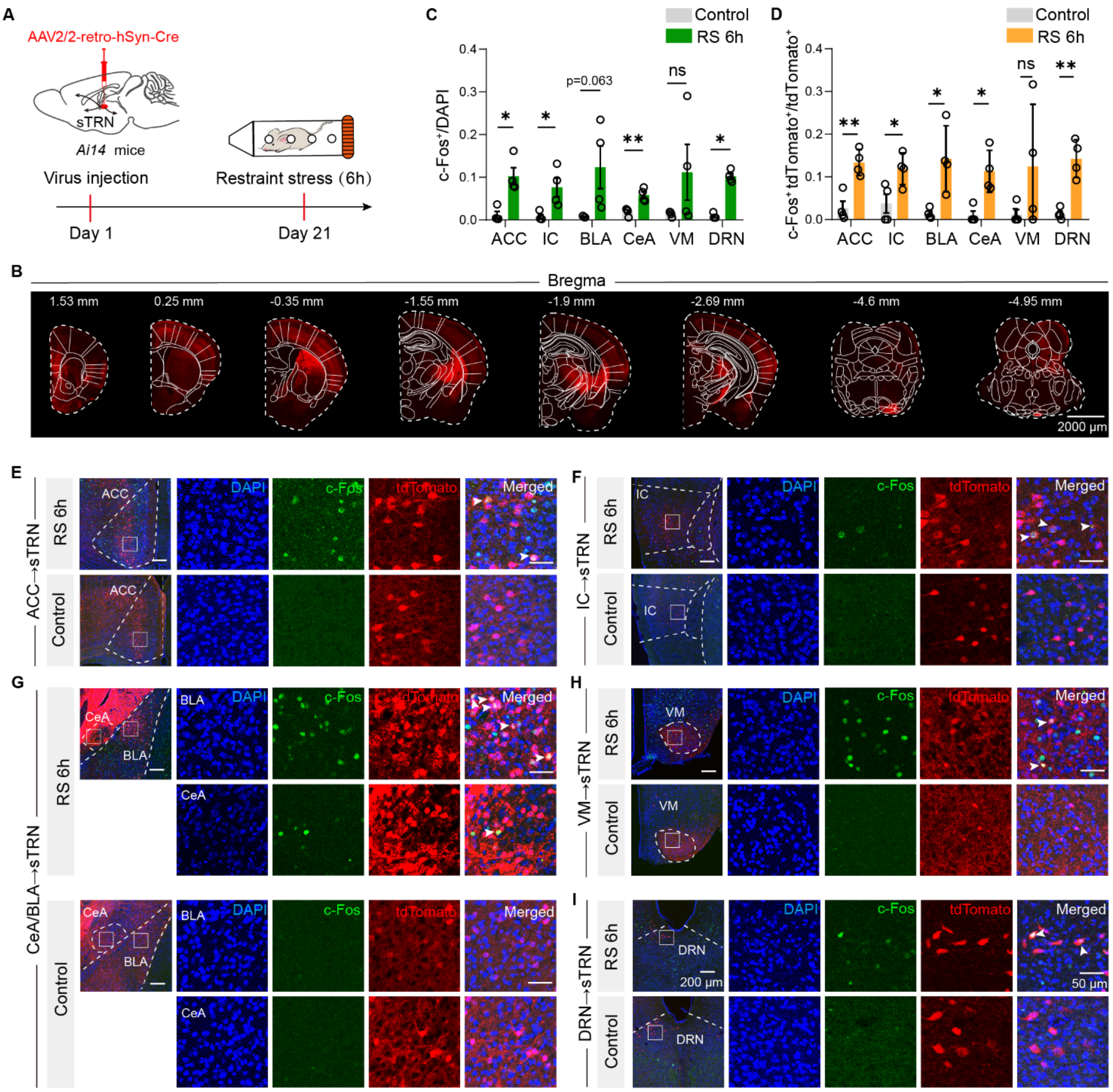


**Fig. S6. sTRN upstream brain regions associated with stress. Related to Fig. 1-Fig. 6.**

(A) Schematic of experimental design.

(B) A series of coronal sections, ipsilateral to the viral injection, from a representative mouse showing brain regions containing sTRN upstream neurons. Scale bar, 2000 μm.

(C) Proportion of c-Fos-labeled neurons in the upstream brain regions of sTRN (n=4 mice/group).

(D) Proportion of tdtomato-labeled neurons expressing c-Fos in upstream brain regions of sTRN (n=4 mice/group).

(E-I) Representative images of retrograde tdtomato$^{+}$ and c-Fos$^{+}$ neurons in ACC (E), IC (F), CeA/BLA (G), VM (H), and DRN (I). Scale bars, 50 μm and 200 μm.

Data are presented as mean ± SEM. “ns”, no significance; $*p < 0.05$, $**p < 0.01$. ACC, IC and DRN of (C), Mann-Whitney U test; others of (C), Unpaired two-sided t test. CeA and VM of (D), Mann-Whitney U test; others of (D), Unpaired two-sided t test.
